# Repetitive transcranial magnetic stimulation targeting motor cortical region induces cell-type-specific transcriptional reprogramming and behavioral recovery in non-human primate model of Rett syndrome

**DOI:** 10.64898/2026.10.20.760437

**Authors:** Wei Li, Wenjie Sun, Jing Su, Futing Yang, Xia Qiu, Yan Zhuo, Zeyuan Yang, Zhengbo Wang, Zhigang Chen, Tianzhuang Huang, Hong Wang, Haoyang Xing, Weizi Ji, Tifei Yuan, Yongchang Chen, Jiaojian Wang

**Author notes:** Correspondence (J.W.), (Y.C.), (T.Y.), (W.J.). These authors contributed equally to this work (Wei Li, Wenjie Sun, Jing Su, Futing Yang, Xia Qiu).

## Abstract

Rett syndrome (RTT), a severe neurodevelopmental disorder caused by mutations in the *MECP2* gene, currently lacks effective treatments. The motor cortex is a pivotal hub in RTT and autism spectrum disorders, and its impairments contribute to motor, social, and cognitive deficits. Here we examined the effects of repetitive transcranial magnetic stimulation (rTMS) targeting motor area in non-human primate of RTT (*MECP2*-mutant monkey) and investigated neuromolecular basis underlying therapeutic effects. We identified a new treatment target, ventral premotor cortex (PMv), for RTT and found that following a 20-day rTMS course, several behavioral and functional connectivity measures showed directional changes. Neuroimaging analyses confirmed that rTMS induced reorganization of large-scale functional connectivity with enhanced functional integration between the somatomotor and the salience networks and normalized coupling between the motor cortex and both the thalamus and the anterior cingulate cortex. Notably, single-nucleus RNA sequencing revealed that rTMS triggered cell-type-specific transcriptional reprogramming: it rescued pathological hyperactivity of excitatory neurons by restoring the expression and regulatory networks of immediate early genes (e.g., *FOS*, *EGR1*) and corrected dysregulated intercellular signaling (e.g., NRG, WNT). These findings indicate that targeting motor cortical areas with brain stimulation may treat RTT, which warrants further clinical validation.

## Introduction

Rett syndrome (RTT) is a neurodevelopmental disorder caused by mutations in the X-linked methyl-CpG-binding protein 2 (*MECP2*) gene (Amir et al. 1999). After a period of apparently normal early development, patients experience progressive regression of motor, cognitive, and social functions, accompanied by stereotyped movements, communication deficits, and autonomic dysfunction (Hagberg et al. 1983). Notably, behavioral studies indicate substantial symptomatic overlap between RTT and autism spectrum disorder (ASD), particularly in social interaction and repetitive behaviors (Neul 2012, Wang et al. 2021). Developing animal models that accurately replicate these clinical manifestations is therefore critical for advancing effective therapies and elucidating the underlying cellular and molecular mechanisms.

The existing RTT models, including *MECP2* mutant mice (Chen et al. 2001, Guy et al. 2001, Patterson et al. 2016, Lombardi, Baker and Zoghbi 2015) and, more recently, non-human primate (NHP) models generated via TALEN-mediated gene editing (Chen et al. 2017), reproduce key behavioral, circuit, and molecular features of the human disease (Wang et al. 2021). Although rodent models have provided fundamental insights into the cellular and synaptic mechanisms underlying RTT pathophysiology and have been widely used to evaluate emerging therapeutic interventions, their neurobiological and phenotypic differences from humans limit translational relevance (Novarino 2017). In contrast, *MECP2*-mutant monkeys exhibit complex behavioral phenotypes and large-scale circuit alterations that overlap with features observed in patients (Chen et al. 2017), providing a complementary model for examining disease-associated phenotypes and evaluating candidate interventions.

Currently, there is no documented effective treatment for RTT. Repetitive transcranial magnetic stimulation (rTMS), a non-invasive neuromodulation technique, has emerged as a promising intervention, capable of targeting dysregulated brain regions to modulate neural activity, restore network homeostasis, and improve behavioral outcomes (Enticott et al. 2014, Sokhadze et al. 2014, Nestor et al. 2022). The primary motor cortex (M1) has been widely reported to be affected in RTT. Neuroimaging studies have reported global brain volume reductions in patients with RTT (Dy-Hollins et al. 2025). At the neurophysiological level, TMS-based assessment of motor cortex has demonstrated that disrupted excitation/inhibition (E/I) balance and the complete abolition of LTP-like plasticity reflect motor disability (Bernardo et al. 2020). Motor-cortex stimulation has also been investigated using transcranial direct current stimulation (tDCS) in RTT and accelerated continuous theta burst stimulation (a-cTBS) in children with ASD (Fabio et al. 2018, Tan et al. 2026), suggesting motor cortex may serve as a biologically grounded and therapeutically actionable target of neuromodulation for RTT.

Here, we examined PMv-targeted rTMS in MECP2-mutant monkeys to characterize behavioral, neuroimaging and transcriptomic associations with cortical stimulation. Multimodal MRI informed target selection, and behavioral and imaging measures were compared before and after intervention where paired observations were available. Single-nucleus RNA sequencing (snRNA-seq) characterized PMv tissue from distinct WT, RTT-pre and RTT-post donor groups. This comprehensive approach not only elucidates the neural plasticity induced by rTMS but also links behavioral recovery to specific cellular and molecular adaptations, thereby providing a mechanistic foundation for precision therapy in Rett syndrome. The overall design of the study is summarized in Figure 1.

**Figure 1.**
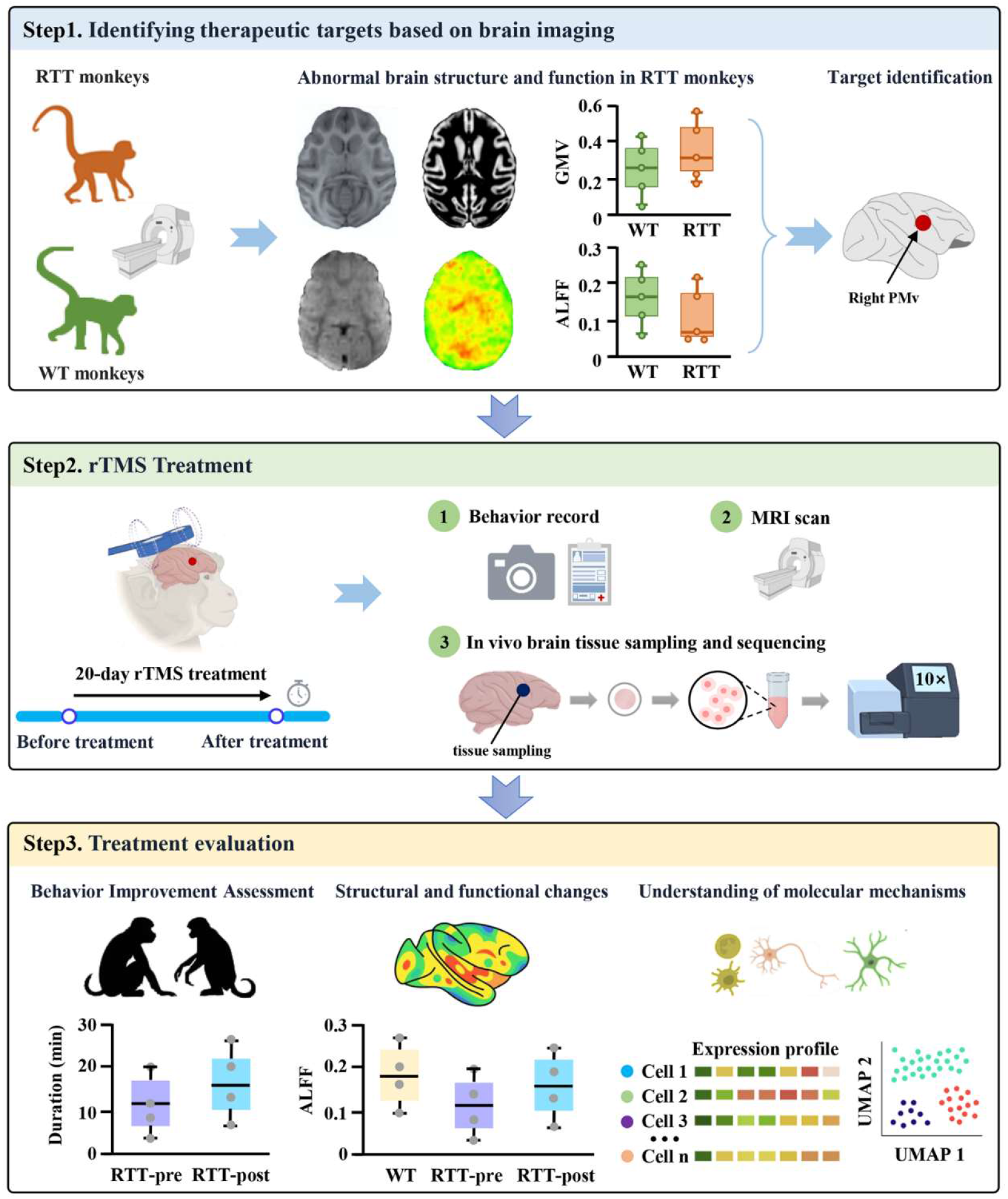
Schematic overview of the experimental design. 1) We firsst acquired structural and functional MRI data from ten monkeys, including five RTT model monkeys and five WT. Structural and functional imaging indices were computed to identify potential cortical targets for intervention, ultimately determining the PMv region as the optimal stimulation site. 2) Based on this target, RTT monkeys underwent a 20-day course of rTMS. Behavioral performance, brain imaging, and single-cell transcriptomic data were collected before and after treatment. 3) Finally, multi-level analyses were performed to evaluate rTMS-induced changes in behavior, neural activity, and molecular profiles at both macroscopic and cellular levels.

## Results

### Identification of rTMS Target Region

To identify a candidate rTMS target, we compared whole-brain resting-state functional and structural MRI measures between *MECP2*-mutant RTT monkeys and age-matched WT controls. RTT animals showed lower amplitude of low-frequency fluctuations (ALFF) in the ventral premotor cortex (PMv; Figure 2a). Structural analysis identified higher grey matter volume (GMV) in the PMv and amygdala and lower GMV in rostral visual area 6 (V6r; Figure 2b). Region-of-interest summaries in the selected PMv region showed lower ALFF (Cohen’s *d* = −8.56, *p* < 0.001) and higher GMV (Cohen’s *d* = 2.04, *p* = 0.032; Figure 2c). The convergence of these differences and the accessibility of PMv to stimulation informed target selection. Following rTMS, ALFF increased (Cohen’s *d* = 3.95, *p* = 0.046) but remained lower than in WT animals (Figure 2c). GMV showed no significant pre/post difference, and the RTT-post versus WT comparison was also non-significant.

**Figure 2.**
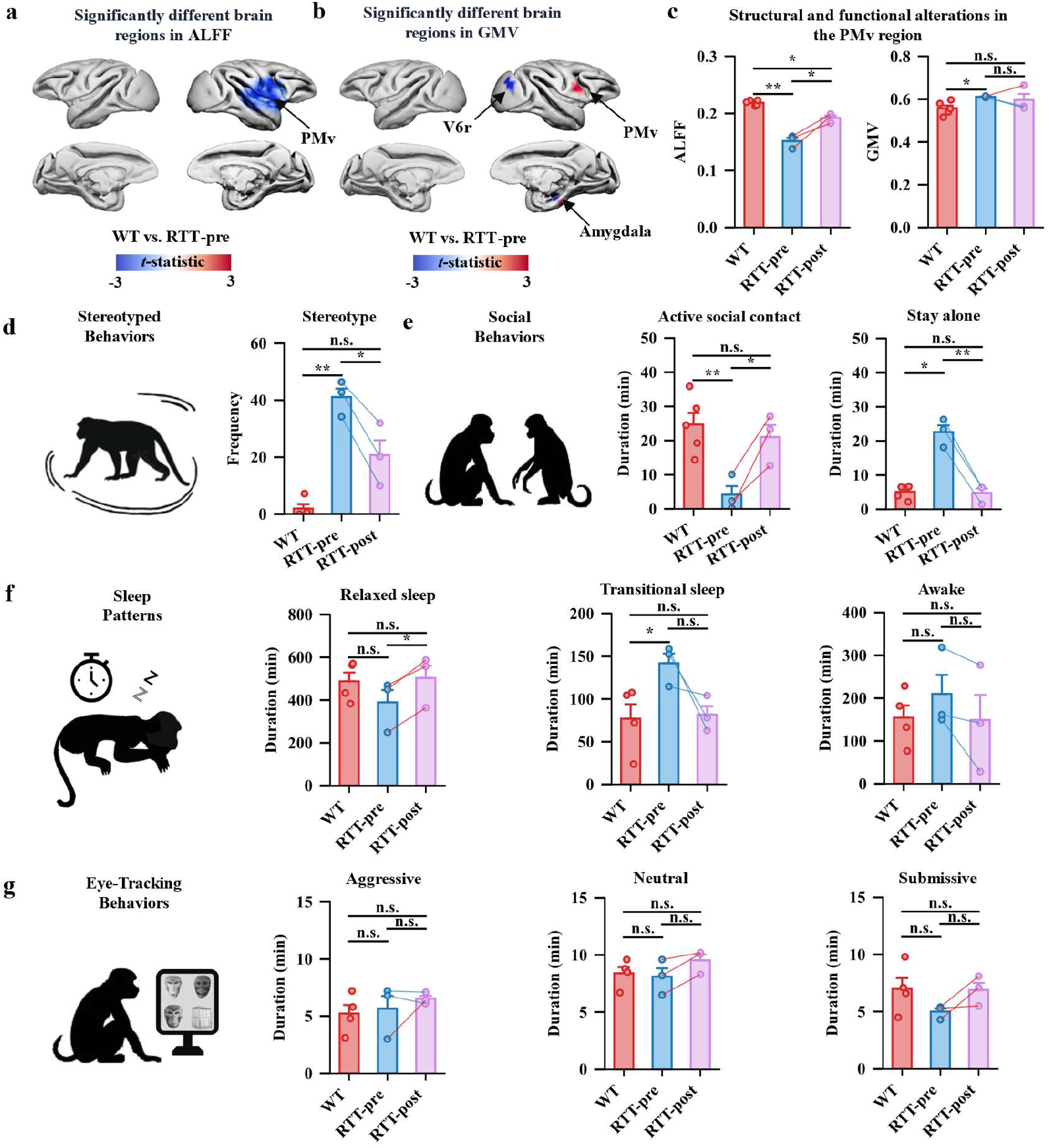
PMv imaging measures and behavioral comparisons before and after rTMS. **(a)** Brain regions showing ALFF differences between RTT-pre and WT animals. **(b)** Brain regions showing GMV differences between RTT-pre and WT animals. **(c)** PMv ALFF and GMV in WT, RTT-pre and RTT-post animals. **(d)** Frequency of stereotyped behavior. **(e)** Duration of active social contact and time spent alone. **(f)** Duration of behaviorally defined relaxed sleep, transitional sleep and wakefulness. **(g)** Fixation duration toward aggressive, neutral and submissive facial expressions. In panels c–*g*, each point represents an individual animal, and lines connect pre-and post-treatment measurements from the same RTT animal. Red lines indicate increases and blue lines indicate decreases. Bars represent mean ± SEM. RTT-pre versus RTT-post comparisons used two-sided paired *t*-tests; comparisons with WT used two-sided Welch’s *t*-tests. \**P* < 0.05, \*\**P* < 0.01, \*\*\**P* < 0.001; n.s., not significant. Abbreviations: PMv, ventral premotor cortex; V6r, rostral visual area 6.

We also examined adult human ASD data to contextualize the monkey imaging findings across a distinct neurodevelopmental diagnosis. The human analysis identified lower PMv ALFF (Cohen’s *d* = −0.38, *p* < 0.001; Figure S1a) and higher GMV (Cohen’s *d* = 0.20, *p* = 0.039; Figure S1b).

### Evaluation of Treatment-Induced Behavioral Changes

Behavioral assessments examined stereotyped behavior, social interaction, sleep and visual attention before and after rTMS. Stereotypy frequency was higher in RTT-pre animals than in WT controls (Cohen’s *d* = 8.15, *p* = 0.002) and decreased after rTMS (Cohen’s *d =* −2.28*, p* = 0.021; Figure 2d). The RTT-post versus WT comparison was non-significant. RTT-pre animals also spent less time in active social contact (Cohen’s *d* = −2.73, *p* = 0.006) and more time alone (Cohen’s *d* = 6.04, *p* = 0.011) than WT controls. After rTMS, active social contact increased (Cohen’s *d* = 2.61, *p* = 0.044), whereas time spent alone decreased (Cohen’s *d* = −5.01, *p* = 0.003; Figure 2e). Neither social measure differed significantly between RTT-post and WT animals.

Behaviorally defined relaxed sleep increased after rTMS (Cohen’s *d* = 0.94, *p* = 0.015), whereas transitional sleep and wakefulness showed no significant pre/post differences (Figure 2f). Transitional sleep was longer in RTT-pre than in WT animals (Cohen’s *d* = 1.93, *p* = 0.042), but its RTT-post versus WT comparison was non-significant. Neither relaxed sleep nor wakefulness differed significantly from WT at either time point. Eye tracking showed no significant pre/post or RTT-versus-WT differences in fixation duration toward aggressive, neutral or submissive facial expressions (Figure 2g).

### Treatment-Induced Alterations in Functional Connectivity

At the network level, functional integration between networks was examined. Following rTMS treatment, enhanced integration between the somatomotor network (SOM) and the salience network (SAL) was observed (RTT-post vs. RTT-pre: Cohen’s *d* = 3.45, *p* = 0.034; Figure 3a). Using the rTMS target region PMv as a seed, whole-brain FC analysis identified two regions exhibiting significant post-treatment changes: the thalamus and the anterior cingulate cortex (CCa) (Figure 3b). Quantitative group comparisons revealed that PMv-thalamic functional connectivity (FC) was significantly elevated in RTT-pre animals relative to WT controls (RTT-pre vs. WT: Cohen’s *d* = 3.17, *p* = 0.004), and was substantially reduced following rTMS intervention (RTT-post vs. RTT-pre: Cohen’s *d* = 6.50, *p* < 0.001; Figure 3c, left). PMv-CCa FC was reduced in RTT-pre animals compared to WT controls (RTT-pre vs. WT: Cohen’s *d* = −1.78, *p* = 0.029), and was restored following treatment in RTT-post animals (RTT-post vs. RTT-pre: Cohen’s *d* = 9.09, *p* = 0.003) (Figure 3c, right), with individual trajectories demonstrating a consistent pattern of recovery across subjects. These findings indicate that rTMS modulation of PMv exerts downstream effects on thalamo-cortical and cingulo-cortical functional coupling.

**Figure 3.**
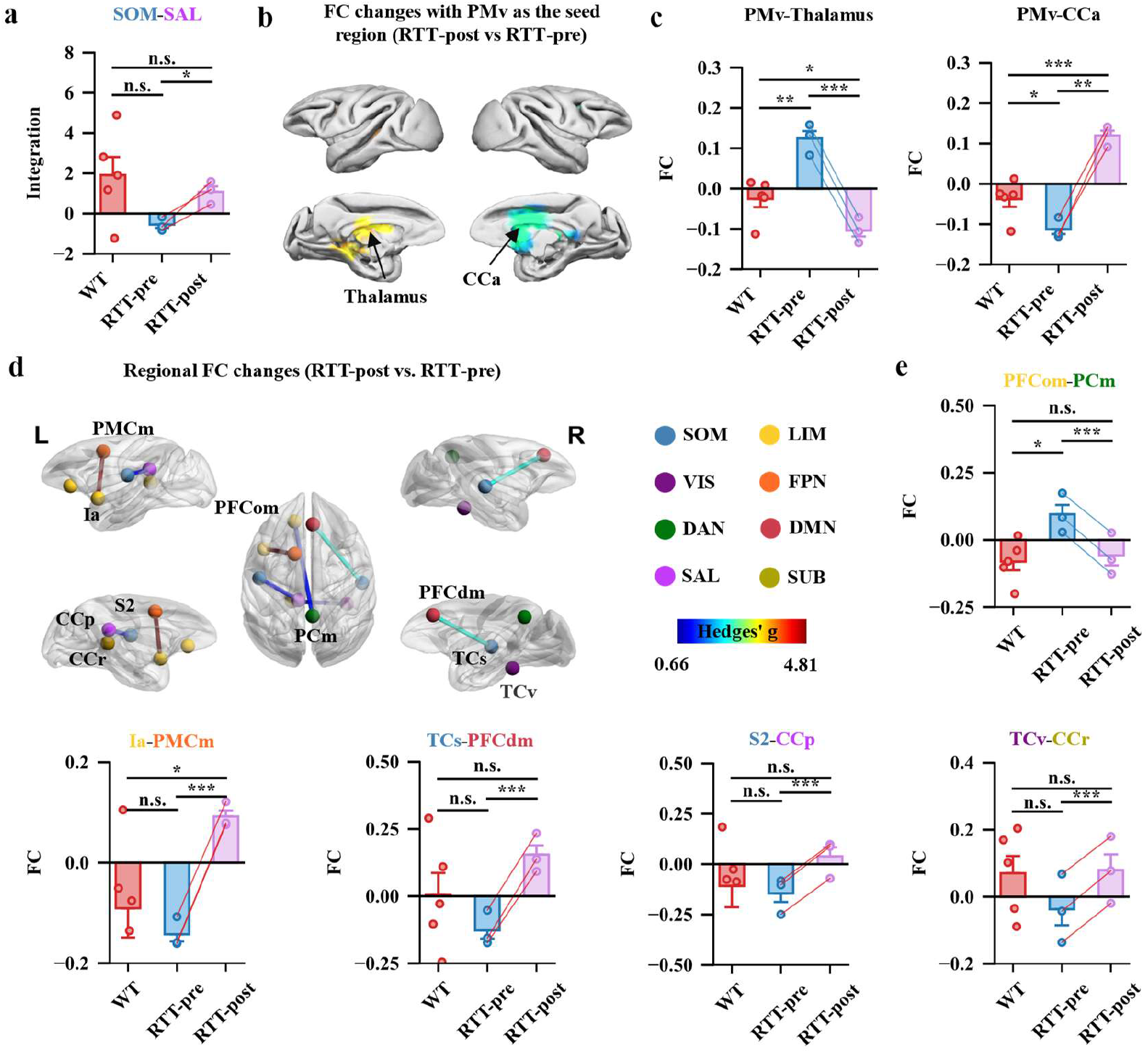
Exploratory functional connectivity comparisons at multiple spatial scales. **(a)** Group comparisons of SOM-SAL functional integration across WT, RTT-pre, and RTT-post groups. **(b)** Whole-brain seed-based FC analysis using PMv as the seed region, selecting two regions at the reported exploratory threshold (RTT-post versus RTT-pre): the thalamus and the CCa, displayed on lateral and medial surface renderings of the macaque brain. **(c)** Quantitative group comparisons of PMv-thalamus FC (left) and PMv-CCa FC (right) across WT, RTT-pre, and RTT-post groups. **(d)** Whole-brain pairwise FC analysis (RTT-post vs. RTT-pre) identifying circuit-level edges exhibiting notable pre/post effect sizes (Hedges’ *g* ranging from 0.66 to 4.81), displayed on lateral and medial surface renderings. Identified connections include PFCom-PCm, Ia-PMCm, TCs-PFCdm, S2-CCp, and TCv-CCr. Color scale reflects Hedges’ *g* magnitude. **(e)** Quantitative group comparisons of individual circuit-level FC pairs across WT, RTT-pre, and RTT-post groups. Lines connecting data points represent individual subjects. Bars represent mean ± SEM. For the animal-level comparisons in panels a, c and e, RTT-pre versus RTT-post comparisons used two-sided paired *t*-tests, and comparisons with WT used two-sided Welch’s *t*-tests. \**P* < 0.05, \*\**P* < 0.01, \*\*\**P* < 0.001; n.s., not significant. Abbreviations: PFCom, orbitomedial prefrontal cortex; PCm, medial parietal cortex; Ia, anterior insula; PMCm, medial premotor cortex; PFCdm, dorsomedial prefrontal cortex; TCs, superior temporal cortex; CCp, posterior cingulate cortex; S2, secondary somatosensory cortex; TCv, ventral temporal cortex; CCr, retrosplenial cingulate cortex; SOM, somatomotor network; VIS, visual network; DAN, dorsal attention network; FPN, frontoparietal network; DMN, default mode network; LIM, limbic network; SUB, subcortical network.

To examine treatment-related reorganization at the level of inter-regional circuits, whole-brain pairwise FC analysis was performed between RTT-post and RTT-pre conditions. Several edges exhibiting notable effect sizes (Hedges’ *g* ranging from 0.66 to 4.81) were identified, including connectivity between prefrontal and parietal regions (PFCom-PCm), between the insula and ventral premotor cortex (Ia-PMCm), between temporal cortex and dorsomedial prefrontal cortex (TCs-PFCdm), between the secondary somatosensory cortex and posterior cingulate cortex (S2-CCp), and between the ventral temporal cortex and retrosplenial cingulate cortex (TCv-CCr) (Figure 3d). PFCom-PCm functional connectivity was found to be significantly elevated relative to WT controls (Cohen’s *d* = 2.26, *p* = 0.027; Figure 3e), whereas rTMS treatment was associated with a subsequent reduction in this connectivity (Cohen’s *d* = 2.03, *p* < 0.001; Figure 3e), suggesting that the modulatory effects induced by rTMS extend beyond the directly stimulated target region to encompass prefrontal-parietal network connectivity. The remaining circuit-level connectivity pairs, including Ia-PMCm, TCs-PFCdm, S2-CCp, and TCv-CCr, exhibited consistent directional increases in functional connectivity following treatment, with individual subject trajectories demonstrating a uniform trend toward normalization.

Exploratory correlations compared pre/post changes in resting-state connectivity with changes in behavioral measures (Figure S2). The displayed pairings included PMv–thalamus connectivity with social contact and sleep duration, and PFCom–PCm connectivity with time alone and neutral-face fixation. Further pairings involved PFCdm–TCs connectivity with wakefulness, and Ia–PMCm connectivity with sleep transitions and dominance-related behavior. These associations are based on a very small number of animals, and their sensitivity to individual observations and multiple testing remains unresolved. They do not establish circuit-specific mediation of behavioral change.

### Single-Cell Genomic Analysis Delineates the Cellular and Molecular Basis of rTMS-Induced Recovery

To systematically delineate the cellular lineages and molecular mechanisms underlying rTMS-mediated functional recovery, single-nucleus RNA sequencing was performed on PMv tissue derived from WT, RTT-pre, and RTT-post monkeys using the 10× Genomics platform. Following stringent quality control via the Seurat pipeline (Stuart et al. 2019), 54,199 high-quality nuclear transcriptomes were retained (median: 1,587 genes and 3,085 transcripts per nucleus) (Figure S3). Unbiased clustering and canonical marker-based annotation identified seven major cell types, OPC, oligodendrocytes, microglia, astrocytes, endothelial cells, excitatory neurons (ExN), and inhibitory neurons (InN), which were further resolved into 21 distinct subtypes (Figure 4a, Figure S4a-b). Pseudobulk differential expression analysis revealed 781 DEGs in RTT-pre vs. WT (microglia exhibiting the highest number) and 1,197 DEGs in RTT-post vs. RTT-pre (L2/3-IT excitatory neurons showing the largest change) (Figure 4b-c, Figure S4c-d). To capture transcriptomic signatures specifically associated with functional recovery, genes exhibiting inverted U-shaped (disease-upregulated, treatment-normalized; *n* = 82, Figure 4d) or U-shaped (disease-downregulated, treatment-restored; *n* = 22; Figure 4e) expression trajectories were defined. Inverted U-shaped genes were predominantly enriched in GABAergic synapse function, negative regulation of cell migration, and axon guidance pathways, whereas U-shaped genes were associated with forebrain development, morphogenesis, and synaptic plasticity signaling (Figure S5).

**Figure 4.**
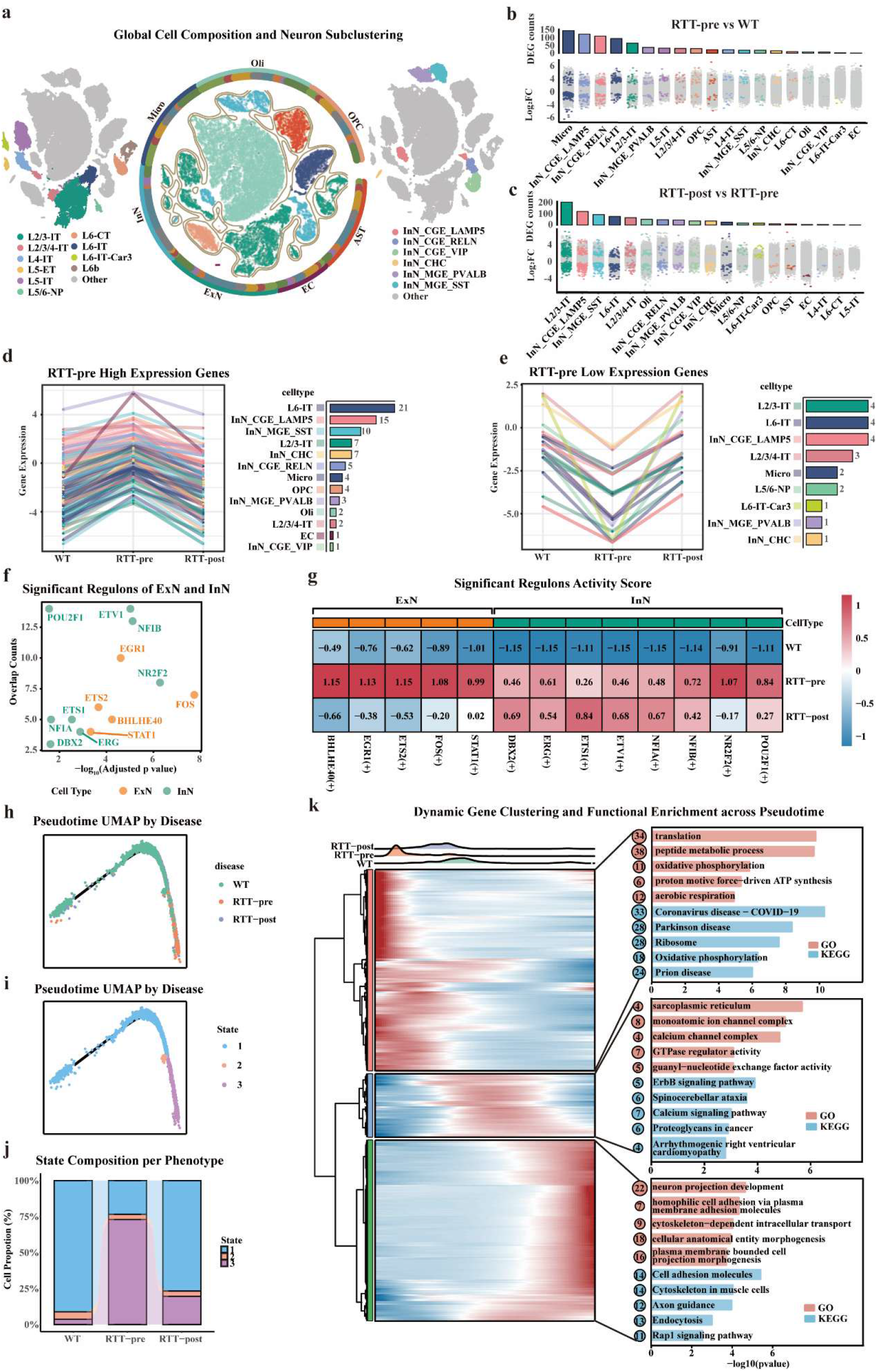
Cross-sectional single-nucleus transcriptomic comparisons among WT, RTT-pre and RTT-post donor groups. **(a)** A total of 54,199 nuclei were retained and classified into seven major cell types and 21 subtypes. The left panel shows the subdivision of excitatory neuronal populations, while the right panel illustrates inhibitory subtypes. Excitatory neurons: L2/3-IT, Layer 2/3 Intratelencephalic neurons; L2/3/4-IT, Layer 2/3/4 Intratelencephalic neurons; L4-IT, Layer 4 Intratelencephalic neurons; L5-ET, Layer 5 Extratelencephalic neurons; L5/6-NP, Layer 5/6 Near-Projecting neurons; L6-CT, Layer 6 Corticothalamic neurons; L6-IT, Layer 6 Intratelencephalic neurons; L6-IT-Car3, Layer 6 Intratelencephalic Car3-expressing neurons; L6b, Layer 6b neurons. Inhibitory neurons: InN_CGE_LAMP5, Inhibitory neuron from Caudal Ganglionic Eminence, LAMP5-positive; InN_CGE_RELN, Inhibitory neuron from CGE, RELN-positive; InN_CGE_VIP, Inhibitory neuron from CGE, VIP-positive; InN_CHC, Chandelier cell; InN_MGE_PVALB, Inhibitory neuron from Medial Ganglionic Eminence, Parvalbumin-positive; InN_MGE_SST, Inhibitory neuron from MGE, Somatostatin-positive. Non-neuronal cells: Micro, Microglia; Oli, Oligodendrocyte; OPC, Oligodendrocyte Precursor Cell; AST, Astrocyte; EC, Endothelial Cell. **(b)** Numbers of DEGs between RTT-pre and WT groups across each cell type. Colored dots indicate significant genes (FDR correction *p* < 0.05). **(c)** Numbers of DEGs between RTT-post and RTT-pre across each cell type. **(d)** Overlapping DEGs between RTT-post vs. RTT-pre and RTT-pre vs. WT that showed higher expression in RTT-pre exhibited an inverted-U pattern (termed inverted U-shaped DEGs, *n* = 82). **(e)** Overlapping DEGs with lower expression in RTT-pre displayed a U-shaped pattern (termed U-shaped DEGs, *n* = 22). **(f)** Overlap analysis between U-shaped and inverted U-shaped DEGs of ExN and InN cell types and the transcriptional regulons identified by SCENIC. Significance was determined using the hypergeometric test, with the y-axis indicating the number of overlapping genes and the x-axis representing the *P*_FDR_ values. **(g)** Regulons with *p*_FDR_ < 0.05 were selected, and their corresponding RAS were examined across different groups to reveal their activity patterns. **(h)** Pseudotime UMAP of microglia, grouped by distinct groups. **(i)** Pseudotime UMAP of microglia, grouped by different states. **(j)** Proportions of microglial cells in each state across different groups. **(k)** Hierarchical clustering of genes significantly associated with pseudotime (1,455 genes identified via GLM fitting with group information) revealed three gene modules: green corresponds to state 1, blue to state 2, and pink to state 3. The distribution of cells along pseudotime across different groups is shown above the heatmap, and the GO and KEGG enrichment results for the three gene modules are displayed on the right.

Exploratory analyses related gene expression to neuroimaging measures and reported five associations at *p* < 0.005 (Figure S6). *NXPH2* expression was negatively correlated with SOM–SAL integration. *COL11A1* and *IGFBP5* expression were negatively correlated with PMv–CCa connectivity, whereas *FAT4* and *SLIT3* were negatively associated with Ia–PMCm and TCv–CCr connectivity, respectively.

### rTMS Restores Excitatory Neuronal Transcriptional Regulatory Networks and Reconfigures Intercellular Communication

To elucidate the transcriptional regulatory architecture underlying rTMS effects, SCENIC-based regulon analysis (Van de Sande et al. 2020) was performed, identifying five significant regulons in ExN and eight in InN that substantially overlapped with U-shaped and inverted U-shaped trajectory genes (Figure 4f, Figure S7). Regulon Activity Score (RAS) comparisons across groups demonstrated that ExN regulons were broadly pathologically upregulated in RTT-pre relative to WT, and were subsequently restored toward baseline levels following treatment, whereas InN regulons remained largely unchanged (Figure 4g). AUCell-based scoring of ExN regulon activity closely recapitulated these RAS patterns, reinforcing the robustness of the identified transcriptional regulatory nodes across independent analytical frameworks (Figure S8). At the intercellular level, L2/3-IT superficial excitatory neurons exhibited pathological hyperactivation of overall communication strength in RTT-pre, which was substantially attenuated following rTMS to levels comparable to WT (Figure S9-S10). Pathway-level dissection identified three signaling cascades, NRG, WNT, and BMP, as particularly salient. NRG-mediated communication from L2/3-IT to inhibitory neurons was aberrantly strengthened in RTT-pre and subsequently suppressed post-treatment; WNT-mediated L2/3-IT outgoing signaling was present in RTT-pre but silenced following rTMS; and BMP-pathway communication between L2/3-IT and L6-IT neurons, absent in RTT-pre, was re-established after treatment (Figure S8b-d). Collectively, these findings position L2/3-IT excitatory neurons as central orchestrators of both transcriptional dysregulation and circuit-level communication remodeling in response to rTMS.

To further contextualize the molecular targets of rTMS within a translational pharmacological framework, L2/3-IT DEGs were cross-referenced against a curated database of 3,547 approved or clinical-stage drugs via ChEMBL (Gaulton et al. 2012) and ToppGene (Chen et al. 2009) enrichment analyses. This integrative approach identified 68 candidate drugs with significant overlap, with DRD2 exhibiting the highest degree of drug connectivity, and Cytarabine, Belinostat, Riluzole, and Amisulpride demonstrating prominent associations with rTMS-responsive gene expression patterns (Figure S11), thereby providing a molecular basis for potential pharmacological synergy with rTMS-based interventions in RTT.

### Microglial Pseudotemporal Dynamics

As the cell type exhibiting the greatest transcriptional divergence between RTT-pre and WT, microglia are likely to play a pivotal role in disease progression. To further dissect the temporal dynamics of microglial states during rTMS treatment, we performed pseudotemporal trajectory analysis on microglial populations, which revealed a continuous transition from homeostatic to disease-associated states, with RTT-pre cells enriched in mid-to-late trajectory states (State 3) and WT and RTT-post cells predominantly occupying the early homeostatic state (State 1), suggesting that rTMS facilitates the reversion of microglia toward a more homeostatic phenotype (Figure 4h-j). GLM-based gene module analysis identified three trajectory-linked modules: the green module (State 1) was enriched in synaptic development, cell adhesion, and axon guidance; the blue module (State 2) was associated with calcium and ErbB signaling; and the pink module (State 3) was characterized by metabolic and bioenergetic pathways including oxidative phosphorylation and ribosomal function (Figure 4k).

## Discussion

In this study, we developed a targeted rTMS protocol focusing on the motor cortex and systematically investigated its therapeutic mechanisms in a primate model of RTT. By integrating multimodal neuroimaging, comprehensive behavioral assessment, and single-nucleus RNA sequencing, we demonstrate that a 20-day 10 Hz PMv-targeted rTMS protocol effectively ameliorates core behavioral deficits in RTT monkeys. At the neural circuit level, rTMS induced widespread functional reorganization, rectifying aberrant connectivity from the motor cortex to the thalamus and anterior cingulate cortex, and modulating the functional coupling between the motor cortex and the salience network. At the molecular level, rTMS normalized pathologically dysregulated transcriptional programs in excitatory neurons, most notably through the restoration of IEG-dependent regulatory networks governed by *FOS* and *EGR1*, and promoted a homeostatic shift in microglial immune states. Furthermore, rTMS reconfigured aberrant intercellular signaling driven by L2/3-IT superficial excitatory neurons, particularly through the NRG, WNT, and BMP pathways. Together, our integrated multiscale analysis provides a systematic characterization of the neural, cellular, and molecular mechanisms underlying rTMS-induced recovery in RTT, and advances its potential as a circuit-precise therapeutic strategy for neurodevelopmental disorders.

Rett syndrome is a neurodevelopmental disorder caused by mutations in the *MECP2* gene, and effective treatments remain limited. Research into its pathophysiology has largely relied on rodent models (Calfa et al. 2011, Na et al. 2012), however, intrinsic neurogenic and phenotypic differences between rodents and humans highlight the need for more translationally relevant models to elucidate disease mechanisms and develop novel therapies (Chen et al. 2017, Novarino 2017). Nonhuman primates, which more closely resemble humans in genetics, brain architecture, cognitive functions, and social behaviors, offer a powerful platform for recapitulating the multilevel pathology of RTT (Aida, Feng and development 2020, Watanabe et al. 2021, Wang et al. 2021). In our previous work, RTT monkeys were shown to exhibit a range of complex behavioral abnormalities including fragmented sleep, increased stereotypies, reduced sensory avoidance, and diminished exploratory behavior, which closely mirror clinical phenotypes observed in patients (Chen et al. 2017). This makes the RTT monkey an ideal model for evaluating neuromodulatory interventions of rTMS which can induce neuronal depolarization, modulate cortical excitability, and enhance synaptic plasticity (Sharbafshaaer et al. 2024, Fitzsimmons et al. 2024).

The selection of PMv as the rTMS stimulation target was grounded in convergent multimodal neuroimaging evidence obtained from both RTT monkeys and human ASD cohorts. The PMv is a key node of the mirror neuron system which is activated both during action execution and action observation (Rizzolatti and Arbib 1998), and plays a central role in action understanding, imitation, empathy, and social cognition (Rizzolatti and Craighero 2004, Gallese, Keysers and Rizzolatti 2004). The 20-day course of rTMS treatment resulted in substantial amelioration across multiple behavioral domains, most notably in stereotyped behaviors and social interaction. The magnitude of improvement in repetitive behavioral repertoires, with post-treatment measures returning to levels statistically indistinguishable from wild-type controls, is particularly noteworthy given the deeply entrenched nature of such behaviors in RTT pathology. Similarly, the partial rescue of social withdrawal, as reflected by increased active social contact and reduced solitary behavior, is consistent with the hypothesis that normalization of motor cortical excitability and connectivity may exert downstream modulatory effects on social motivational circuits (Iacoboni and Dapretto 2006, Kang et al. 2025).

The present findings demonstrate that PMv-targeted rTMS induces widespread functional reorganization across multiple levels of neural circuitry in RTT, encompassing thalamocortical, prefrontal-parietal, and large-scale inter-network connectivity. The pathological hyperconnectivity observed between PMv and the thalamus in untreated RTT animals is consistent with established evidence that *MECP2* loss-of-function disrupts cortical excitatory-inhibitory balance through impaired GABAergic interneuron maturation (Li 2022, Liiwand et al. 2026), thereby destabilizing thalamocortical feedback regulation. The substantial normalization of PMv-thalamic coupling following rTMS intervention, alongside the reciprocal restoration of PMv-CCa connectivity, suggests that non-invasive premotor stimulation is capable of rectifying circuit-level dysregulation through mechanisms that extend beyond the directly targeted region, a finding in line with prior evidence that rTMS-induced plasticity propagates via polysynaptic corticothalamic pathways (Hallett 2007, Nettekoven et al. 2015). At the network level, widespread changes in functional connectivity were observed between multiple regions of the motor cortex and the SAL and DMN networks, characterized by a directional reduction in TCs-PFCdm coupling and a synchronous increase in S2-CCP connectivity. Furthermore, following rTMS treatment, enhanced integration from the SOM to SAL was observed. The SOM-SAL integration was reduced in RTT-pre relative to WT controls, indicating that disrupted somatomotor-salience coupling may constitute a pathological network signature of RTT. The salience network plays a pivotal role in filtering interoceptive and sensory signals (Uddin 2015), and its functional dysconnectivity has been consistently reported in autism spectrum disorder and related neurodevelopmental conditions (Attanasio et al. 2025). We found that SOM-SAL integration was negatively correlated with *NXPH2*, a synaptic gene, suggesting that motor cortex rTMS may modulate large-scale network organization via synaptic mechanisms (Wilson et al. 2019, Missler and Sudhof 1998). The concurrent normalization of multiple inter-regional connectivity pairs, including PFCom-PCm, TCv-CCr, and Ia-PMCm, further indicates that rTMS modulation of a single premotor hub exerts distributed reorganizational effects across functionally distinct circuits subserving social cognition, sensorimotor integration, and sleep-wake regulation. Critically, the systematic correlations between connectivity changes and behavioral recovery across multiple domains provide translational validation for the circuit-level findings, suggesting that the therapeutic efficacy of rTMS is mechanistically coupled to the functional re-integration of specific, identifiable neural pathways rather than reflecting a non-specific neuromodulatory effect (Hallett 2007, Kang et al. 2025).

At the transcriptomic level, we found that the transcriptional changes in excitatory neurons appear to be mediated by immediate-early genes (IEGs), notably *FOS* and *EGR1*, whose regulatory networks were pathologically activated in RTT but subsequently suppressed following rTMS treatment. IEGs are rapidly and transiently induced by neuronal activity and typically encode transcription factors that orchestrate downstream gene expression, thereby influencing synaptic plasticity, learning, and memory (Pérez-Cadahía et al. 2011, Morgan and Curran 1989, Morgan and Curran 1991, Marballi and Gallitano 2018). Dysregulation of IEGs has been reported in RTT (Kron et al. 2012, Swanberg et al. 2009, Li et al. 2013), and studies in rodents show that rTMS robustly induces *FOS* and *EGR1* in cortical neurons (Aydin-Abidin et al. 2008, Fujiki, Yee and Steward 2020). Our findings extend this observation to a non-human primate disease model, providing cross-species evidence that rTMS selectively activates these excitatory transcription factors. In contrast, the regulatory networks of inhibitory neurons did not show significant changes in activity after rTMS. Although certain inhibitory neuron-associated transcription factors exhibited differential expression, these alterations did not translate into broad shifts in regulon activity, suggesting that the rTMS protocol used here may not substantially reprogram the transcriptional landscape of inhibitory neurons.

At the circuit level, cell-cell communication analysis revealed that L2/3-IT neurons exhibited pathological hyperactivation in their outgoing signaling reflecting excessive excitatory drive within cortical networks. rTMS significantly attenuated this aberrant communication intensity restoring it toward wild-type levels, which is consistent with reported rTMS-induced plasticity in layer 2/3 neurons (Tang et al. 2021). We focused on three signaling pathways linked to RTT pathology: NRG, WNT, and BMP. NRG signaling which regulates axon guidance, synapse formation, and excitatory-inhibitory balance (Falls 2003, Mei and Xiong 2008, Mei and Nave 2014) was overactivated in RTT and normalized by rTMS. Similarly, dysregulated WNT signaling, implicated in cognitive development and previously shown to be modulated in RTT mouse models (Kwan, Unda and Singh 2016, Hsu et al. 2020), was restored by rTMS rebalancing communication from L2/3-IT neurons to other populations. Finally, BMP signaling, whose genes are closely tied to *MECP2* deficiency, cortical development, dendritic morphogenesis, synapse formation, and glial cell differentiation (Hong et al. 2023, Nakashima et al. 2021), was also reconstituted by rTMS, particularly in communication between L2/3-IT and L6-IT neurons. Taken together, these findings indicate that aberrant BMP, WNT, and NRG signaling contribute to disrupted local circuits in RTT, and their coordinated normalization by rTMS may facilitate the recovery of neuronal plasticity.

Neuroimmune dysregulation is increasingly recognized as a key component of RTT pathophysiology (Derecki, Privman and Kipnis 2010, Jin et al. 2015). Microglia, the central nervous system resident immune cells, are essential for synaptic refinement, homeostasis, and response to neuronal stress (Salter and Stevens 2017). Consistent with prior studies (Zhao et al. 2017, Schafer et al. 2016), microglia were identified as the most transcriptionally altered cell type in RTT-pre relative to WT. Pseudotemporal trajectory analysis revealed a continuous transition from homeostatic to disease-associated microglial states, with RTT-pre cells enriched in mid-to-late trajectory states (State 3), whereas WT and RTT-post cells were predominantly distributed in the early homeostatic state (State 1), suggesting that rTMS facilitates the reversion of microglia toward a more homeostatic phenotype. Functional enrichment along the trajectory delineated three microglial states: a homeostatic State 1 enriched in WT and rTMS-treated samples and associated with cell adhesion, synaptic development, and axon guidance; a disease associated State 3 predominant in untreated RTT and characterized by elevated oxidative phosphorylation and ribosomal activity, reflecting chronic metabolic activation (Depp et al. 2025, Färber and Kettenmann 2006); and an intermediate State 2 with heightened calcium and ErbB signaling. rTMS redirected microglial from the activated State 3 back toward the homeostatic State 1. These findings position microglia as both sensors and effectors of cortical remodeling following rTMS-based neuromodulation (Hanisch and Kettenmann 2007), and raise the possibility that dampening of pathological microglial activation constitutes an important cellular mechanism contributing to functional recovery in RTT.

The cross-species validation of PMv abnormalities, across both *MECP2*-mutant primate and human ASD cohorts, and the convergence of behavioral, neuroimaging, and transcriptomic evidence collectively strengthen the translational relevance of the present findings and support the development of motor cortex targeted rTMS protocols for clinical investigation in RTT and ASD patients. Nevertheless, several important limitations warrant acknowledgment. The small sample sizes inherent to non-human primate studies constrain statistical power, particularly for the sleep and affective processing domains in which only trend-level effects were observed. The causal directionality of the identified brain-behavior correlations remains to be established through circuit-specific perturbation approaches, and the generalizability of the present stimulation parameters to other rTMS protocols and cortical targets remains to be systematically evaluated. Future studies employing larger cohorts, longitudinal follow-up designs, and mechanistic circuit interrogation will be essential to consolidate and extend the present findings toward clinical translation.

In summary, this study provides an integrated neurobiological framework for rTMS-induced recovery in a non-human primate model of RTT, spanning behavioral, circuit, cellular, and molecular levels of analysis. The convergent evidence demonstrates that PMv-targeted rTMS ameliorates core RTT deficits through the coordinated normalization of thalamocortical connectivity and the enhancement of functional integration between the motor cortex and the salience network, the restoration of IEG-driven transcriptional regulatory programs in excitatory neurons, the reconfiguration of intercellular signaling networks dominated by superficial L2/3-IT neurons, and the promotion of homeostatic microglial states. These findings collectively position motor cortical stimulation as a mechanistically grounded and translationally promising therapeutic strategy for RTT and related neurodevelopmental disorders, warranting rigorous validation in clinical settings.

## Materials and Methods

### Animals and Subject Acquisition

This study aimed to investigate the neurobiological basis of rTMS intervention in monkeys with RTT. Experimental animals were obtained from the State Key Laboratory of Primate Biomedical Research at Kunming University of Science and Technology. A total of 10 female cynomolgus monkeys including 5 wild-type controls (WT; age = 8.8 ± 0.84 years) and 5 RTT monkeys (age = 9.0 ± 0.82 years) were enrolled. The RTT monkeys were generated via TALEN-mediated mutation of the *MECP2* gene and, as demonstrated in our previous studies, exhibit behavioral phenotypes (Chen et al. 2017, Wang et al. 2021) and functional connectivity dynamics (Li et al. 2025) that closely resemble those observed in patients. During the experimental period, one RTT monkey died due to natural causes and was therefore excluded from post-treatment behavioral and imaging analyses. However, this animal had completed brain tissue acquisition after death for further single-cell RNA sequencing. All experimental procedures strictly adhered to national guidelines for animal care and use and were approved by the relevant authorities in China and the Institutional Animal Care and Use Committee of Kunming University of Science and Technology (Permit No.: KUST202401002).

### Monkey MRI Data Acquisition

To comprehensively evaluate the structural and functional brain alterations in RTT monkeys pre-and post-intervention, all subjects underwent high-resolution T1-weighted structural MRI and resting-state functional MRI (rs-fMRI) on a 3.0 T Siemens Prisma scanner equipped with a dedicated monkey head coil. Animals were anesthetized via intramuscular injection of Zoletil (0.1 mL/kg) and Sumianxin (0.1 mL/kg), achieving adequate sedation and muscle relaxation. Anesthesia was maintained for approximately 1 hour, during which heart rate, respiration, and body temperature were continuously monitored to ensure physiological stability and optimal image quality.

Structural images were acquired using a 3D magnetization-prepared rapid gradient-echo (3D-MPRAGE) sequence with TR = 1400 ms, TE = 3.55 ms, TI = 900 ms, flip angle = 8°, FOV = 135 × 135 mm², matrix = 224 × 224, slice thickness = 0.6 mm, voxel resolution = 0.6 × 0.6 × 0.6 mm³, 160 slices, acceleration factor = 2, and NEX = 3. The functional images were obtained using a gradient echo-planar imaging (EPI) sequence (TR = 2000 ms, TE = 22 ms, flip angle = 90°, matrix = 64 × 64, voxel resolution = 1.5 × 1.5 × 1.5 mm³, 250 volumes). During rs-fMRI acquisition, monkeys were maintained in a resting, eyes-closed state to minimize sensory input and ensure functional signal fidelity.

### Preprocessing of Structural and Functional MRI Data of Monkeys

Structural MRI was first preprocessed including resampling, removal of non-brain tissues, intensity normalization, and bias field correction. Individual T1-weighted images were then registered to the standard monkey brain template (INIA19) (Rohlfing et al. 2012), followed by tissue segmentation using a probabilistic atlas to generate gray matter (GM), white matter (WM), and cerebrospinal fluid (CSF) probability maps. Gray matter probability maps were subsequently smoothed with a 3 mm full-width at half-maximum (FWHM) Gaussian kernel to enhance the signal-to-noise ratio, and individual gray matter volume (GMV) was computed for subsequent statistical analysis.

Functional MRI preprocessing involved discarding the first 10 time points to achieve magnetization equilibrium, followed by slice-timing correction and head motion correction. All volumes were realigned to the first time point to minimize residual motion artifacts. Functional images were then co-registered to their corresponding structural images and normalized to the template space. Spatial smoothing was applied using a 3 mm FWHM Gaussian kernel, and temporal filtering (0.01-0.05 Hz) was performed to retain low-frequency fluctuations. To reduce non-neuronal noise, head motion parameters, as well as WM and CSF signals, were regressed out. The ALFF was calculated for each individual to reveal abnormal functional activity in RTT monkeys.

### Definition of rTMS Target Brain Region in Monkey

In this study, we used the brain region showing both structural and functional alterations in RTT monkeys as the therapeutic target. To identify structural alterations, a voxel-wise two-sample t-test was conducted between RTT monkeys before treatment and WT monkeys to identify abnormal GMV, with multiple comparisons corrected across the whole brain using Gaussian random field (GRF) theory (thresholded at *p* < 0.05). To further reveal abnormal functional activity, the two-sample t-test was also conducted to identify ALFF differences between RTT and WT monkeys, and the significance level was determined using GRF correction method at *p* < 0.05. Brain region showing significant differences in both GMV and ALFF was considered as treatment target for exploring therapeutic mechanisms and guiding single-cell sampling analyses.

### Comparison with human ASD imaging data

RTT patients often exhibit ASD-like symptoms during early development, including social communication deficits and stereotyped behaviors (Neul 2012, Percy 2011). RTT monkeys have been widely used to investigate ASD mechanisms, as they partially recapitulate the molecular and circuit-level underpinnings of the disorder (Wu et al. 2021, Chahrour and Zoghbi 2007). Leveraging Autism Brain Imaging Data Exchange (ABIDE) as a large-scale, publicly accessible neuroimaging resource (Di Martino et al. 2014, Di Martino et al. 2017), we selected 200 adult human ASD participants (age = 26.69 ± 10.15 years; male/female = 189/11) whose ages were matched to those of the RTT monkeys, along with 258 healthy controls (age = 27.02 ± 9.34 years; male/female = 232/26). Detailed scanning parameters and demographic information are provided in Supplementary Tables S1 and S2.

For the preprocessing of human structural and functional MRI data, we employed the fMRIPrep (v22.1.1) pipeline (Esteban et al. 2019), which automatically generates a standardized and reproducible workflow (details are provided in the Supplementary Methods). Briefly, structural (T1-weighted) images were first corrected for intensity nonuniformity (Tustison et al. 2010), followed by skull stripping (Avants et al. 2008). The brain was then segmented into CSF, WM, and GM, and cortical surfaces were reconstructed (Dale, Fischl and Sereno 1999). Finally, individual structural images were nonlinearly registered to both the *MNI152NLin6Asym* and *MNI152NLin2009cAsym* template spaces (Evans et al. 2012). For statistical analyses, voxel-wise gray matter probability maps derived from structural segmentation were smoothed with an 8 mm FWHM Gaussian kernel.

For functional images, preprocessing began with motion correction (Jenkinson et al. 2002) and slice-timing correction (Cox, Hyde and Vivo 1997), followed by boundary-based co-registration of the BOLD data to each subject’s T1-weighted reference image (Greve and Fischl 2009). From the preprocessed BOLD data, we computed a series of nuisance regressors, including head motion parameters (FD, DVARS) (Power et al. 2014), three global signals (CSF, WM, and whole-brain) (Jenkinson et al. 2002), and noise principal components (Behzadi et al. 2007). To enhance denoising performance, first-order derivatives and quadratic terms were included in the confound regression model. Frames exceeding FD > 0.5 mm or DVARS > 1.5 were identified and flagged as motion outliers. Subsequently, the BOLD data were normalized to the *MNI152NLin6Asym* space and further denoised using ICA-AROMA (non-aggressive mode) to mitigate motion-related artifacts (Pruim et al. 2015). Spatial smoothing was then applied using a 6 mm FWHM Gaussian kernel. Finally, the residual time series were obtained by regressing out motion parameters, WM, CSF, and global signals, followed by band-pass filtering within the range of 0.01-0.1 Hz. ALFF was calculated for each participant to characterize functional activity. To account for multi-site scanning variability, individual ALFF maps were standardized prior to group comparisons.

To identify structural and functional abnormalities, two-sample *t*-tests on GMV and ALFF were conducted, controlling for age, sex, head motion and scanning site. The multiple comparisons were corrected using GRF theory, with a significance threshold of *p* < 0.05.

### rTMS Intervention Protocol

After determining the treatment target brain region showing both GMV and ALFF alterations in the above analyses, high-frequency rTMS was applied to RTT monkeys to improve brain functional activity. Stimulation parameters were as follows: a 10-Hz frequency, a 3.3 s stimulation duration with 17 s inter-train intervals, at 80% of the individual motor threshold (MT). This sub-threshold intensity was deliberately selected to minimize the risk of seizure induction. Each session delivered a total of 2,079 pulses over 21 minutes. The intervention spanned 20 days, with the first 10 treatments conducted consecutively, followed by a 1-day break, and the remaining 10 treatments thereafter. To ensure precise targeting and animal immobilization, a custom high-density ethylene-vinyl acetate (EVA) cap was used to expose the stimulation site and provide stable restraint.

### Stereotyped Behaviors

Stereotyped motor behaviors were assessed via continuous 70-min video recording in a standardized monkey cage under controlled lighting and minimal external disturbance. Two primary behavioral metrics were quantified: the total duration (in seconds) and frequency (number of occurrences) of circling behavior, defined as repetitive rotational locomotion exceeding 360° in a consistent direction. Locomotor activity trajectories were simultaneously captured using automated tracking software [PrimateScan system (Clever Sys Inc.)], which generated spatial heatmaps and path length measurements to provide an objective index of stereotypy.

### Social Behaviors

Social interaction was assessed using a validated “social cage” paradigm employing a dual-chamber apparatus separated by a removable opaque partition. Each subject underwent a 15-min isolation period in one chamber to establish a social motivation baseline, after which the partition was removed to allow 45 min of spontaneous free interaction with a conspecific. The following behavioral variables were quantified: the frequency of proactive social contact initiations and the cumulative duration of social interaction. Non-social behaviors, including self-directed behaviors and solitary locomotion, were also recorded to provide contextual comparison.

### Eye-Tracking Behaviors

Eye tracking was employed as a non-invasive approach to assess cognitive and social information processing in RTT monkeys, using a Tobii Pro TX300 Eye Tracker (Tobii Technology AB, Danderyd, Sweden). Gaze data were sampled at 300 Hz and displayed on a 23-inch monitor (resolution: 1920 × 1080 pixels) positioned approximately 69 cm from the subjects’ eyes. Prior to testing, all monkeys underwent a habituation protocol in which they were trained to sit quietly in a primate chair with head movement restricted by a customized high-density ethylene-vinyl acetate (EVA) helmet fitted to each individual. Two consecutive paradigms were administered with an approximately one-week interval: (1) a paired face-object preference test and (2) a paired emotional faces preference test. Prior to each session, a two-point calibration procedure was performed using Tobii Studio software, with animated audio symbols employed as attentional cues. Ambient lighting was reduced throughout testing to minimize distractions. In the paired face-object preference test, 20 stimuli were presented, each comprising a conspecific neutral face paired with a familiar object. The stimulus set included faces from 7 individuals (10 faces total) paired with 10 distinct objects, arranged with counterbalanced lateralization. Face and object stimuli subtended approximately 8.5 × 9° of visual angle and were separated by 8.5°. In the paired emotional faces preference test, 14 stimuli were presented, each consisting of two facial expressions from the same individual. Stimuli were drawn from 5 monkeys, each displaying two or three expressions, encompassing three emotional contrast conditions: neutral-aggression, neutral-submission, and aggression-submission, presented in counterbalanced sequence. All monkey facial stimuli were sourced from the PrimFace database (https://visiome.neuroinf.jp/primface). For both paradigms, the left and right halves of each stimulus display, demarcated by the screen midline, were defined as two areas of interest (AOIs) corresponding to the respective face or object. Looking frequency and cumulative looking duration within each AOI were automatically extracted using Tobii Studio software (Tobii Technology AB, Danderyd, Sweden).

### Sleep Patterns

Sleep-wake architecture was evaluated through continuous 12-h nocturnal video recording (20:00-08:00) in a specialized sleep-monitoring cage under dim red lighting to preserve natural sleep behavior while maintaining recording visibility. Behavioral sleep states were classified into three categories: wakefulness, transitional sleep (characterized by recumbent posture with intermittent movement and partial eye closure), and relaxed sleep (characterized by sustained recumbency, absence of spontaneous movement, and complete eye closure). The proportional duration of each sleep state relative to the total recording period was calculated for each subject.

### Functional Connectivity Analysis

To evaluate the effects of rTMS on PMv-centered functional connectivity, a seed-based whole-brain FC analysis was performed using the PMv as the seed region. Specifically, the mean blood oxygen level-dependent (BOLD) time series was extracted from the PMv, and Pearson correlation coefficients were computed between the seed time series and the time series of every voxel in the brain, generating individual whole-brain functional connectivity maps. The resulting correlation coefficients were then transformed into z-scores using Fisher’s r-to-z transformation for subsequent group-level statistical analysis. Paired t-tests were conducted to compare differences between the RTT-pre and RTT-post conditions. Clusters exceeding 30 voxels with *p* < 0.05 were identified and retained for subsequent between-group comparisons.

At the level of circuit-level reorganization, whole-brain ROI-to-ROI pairwise functional connectivity analysis was conducted based on the D99 macaque brain atlas (Reveley et al. 2017), generating a complete FC matrix. After Fisher z-transformation, paired t-tests were applied to identify group differences, and connections with notable effect sizes were selected as candidate treatment-responsive edges.

At the large-scale network level, brain regions were assigned to eight canonical functional networks: somatomotor network (SOM), visual network (VIS), dorsal attention network (DAN), salience/ventral attention network (SAL), frontoparietal network (FPN), default mode network (DMN), limbic network (LIM), subcortical network (SUB). To further quantify coupling between the eight canonical networks, we adopted the network allegiance framework adapted from Daws et al. (Daws et al. 2022). For each subject, the Louvain community detection algorithm was applied to the functional connectivity matrix across 100 iterations. A region-by-region allegiance matrix P (size *N* × *N*) was constructed by computing, for each pair of regions, the proportion of iterations in which the two regions were assigned to the same module. For each of the 28 unique pairs among the eight canonical networks, an integration index was defined as the mean allegiance across all inter-network region pairs connecting the two networks. To control for differences in network size and chance-level co-assignment, allegiance values were normalized against a null distribution generated by randomly permuting region-to-network labels 1,000 times, yielding a z-scored integration index for each network pair and subject.

### Samples Acquisition for snRNA-seq

To investigate the molecular and cellular mechanisms underlying rTMS effects, in vivo single-nucleus sampling was performed in the stimulation target region identified through MRI analysis. A total of five RTT monkeys were included in this procedure. One monkey, together with the deceased monkey, was classified into the RTT-pre group. The remaining three monkeys underwent tissue collection following completion of treatment and were designated as the treatment group for characterization of the therapeutic effects of rTMS. Three randomly selected WT monkeys served as controls. All tissue samples were snap-frozen in liquid nitrogen and stored at-80°C to prevent batch effects and were processed simultaneously. Tissue samples were processed for library construction using the 10x Genomics Chromium platform.

### Isolation of nuclei and single-nucleus RNA sequencing

PMv tissue samples from WT, untreated RTT (RTT-pre), and rTMS-treated RTT (RTT-post) monkeys were processed in parallel within the same nuclei-isolation batch to limit technical variation. Frozen samples were kept on dry ice until processing and were homogenized in ice-cold RNase-free lysis buffer containing 0.32 M sucrose, 5 mM CaCl₂, 3 mM MgAc₂, 0.1 mM EDTA, 10 mM Tris-HCl, 1 mM DTT, and 0.1% Triton X-100 in DEPC-treated water. Homogenization was performed using a glass Dounce homogenizer until no visible tissue fragments remained. The homogenates were briefly incubated on ice to promote nuclear release and then passed through a 30-μm strainer to remove tissue debris. Following centrifugation, nuclei were further enriched by density-gradient ultracentrifugation. The purified nuclear pellet was resuspended in DEPC-treated PBS, maintained briefly on ice, and filtered twice through 30-μm strainers to eliminate residual aggregates. Nuclear concentration and morphology were evaluated using light microscopy and hemocytometer counting. Only intact, translucent nuclei with minimal aggregation and background debris were used for library construction. Qualified nuclei were adjusted to a concentration of approximately 2,000 nuclei/μL and loaded onto the 10x Genomics Chromium platform according to the manufacturer’s protocol. Libraries generated from individual samples were pooled and sequenced on an Illumina NovaSeq platform, targeting approximately 50,000–100,000 reads per nucleus.

### Genomics Data Pre-processing and Annotation

Sequencing data in FASTQ format were processed using Cellranger (v7.1.0) for alignment, filtering, barcode counting, and UMI counting, with the Mmul_10 Macaca mulatta genome (Zheng et al. 2017). Subsequent analysis of the count expression matrix was performed using the Seurat package (v5.1.0) in R (Stuart et al. 2019), and the following filtering criteria were applied to exclude low quality cells: fewer than 500 detected genes, fewer than 1000 UMIs, mitochondrial gene content exceeding 5%, and ribosomal gene content exceeding 5%. In addition, potential ambient RNA contamination was assessed using decontX package (v1.2.0) (Yang et al. 2020), and cells with a contamination rate greater than 0.2 were removed. Putative doublets were identified and excluded using the DoubletFinder package (v2.0.4) (McGinnis, Murrow and Gartner 2019), based on clustering of major cell populations. After filtering, high-quality cells were retained for downstream analysis. The dataset was stratified by sample origin, and the raw count normalization was applied using the NormalizeData function with a scale factor of 10,000, followed by identification of the top 2,000 highly variable genes (HVGs) using the FindVariableFeatures function. To mitigate batch effects across samples, we applied the Harmony algorithm to integrate the datasets. Subsequently, principal component analysis (PCA) was performed on the integrated data, after which the JackStraw algorithm and ElbowPlot were employed to determine significant principal components (PCs). The top 23 PCs were selected for downstream analyses, including k-nearest neighbor (KNN) graph construction and clustering via the shared nearest neighbor (SNN) algorithm. For visualization, uniform manifold approximation and projection (UMAP) embeddings were generated in two dimensions. Based on canonical marker genes, a total of 21 distinct clusters were annotated.

### Differential Gene Expression and Molecular Enrichment

Differential gene expression analysis across distinct cell types was performed using the pseudo-bulk approach with the likelihood ratio test (LRT) implemented in the edgeR framework via Libra package (v 1.0.0) (https://github.com/neurorestore/Libra.git) to identify genes differentially expressed between WT and RTT-pre, and between RTT-pre and RTT-post. Because all animals were female and age did not differ significantly among groups, no additional demographic covariates were included in the differential-expression models. Genes with a log_2_ fold change of expression of at least ± 0.25 and Benjamini-Hochberg FDR *P* < 0.05 were considered statistically significant. Subsequently, U-shaped and inverted U-shaped trajectory genes were defined as those that were significantly differentially expressed in both the RTT-pre vs. WT and RTT-post vs. RTT-pre comparisons within the same cell type. Here, “U-shaped” denotes genes with low expression in RTT-pre, whereas “inverted U-shaped” refers to those with high expression in RTT-pre. Gene Ontology (GO) and Kyoto Encyclopedia of Genes and Genomes (KEGG) pathway enrichment analyses were then performed on these defined gene sets using the clusterProfiler package (v 4.12.6) (Yu et al. 2012). These analyses were performed to elucidate the predominant molecular functions and biological pathways associated with the observed transcriptional changes.

### Drug-target enrichment analysis

Given that L2/3-IT neurons exhibited the largest number of differentially expressed genes between the pre-and post-treatment conditions, we subsequently leveraged two comprehensive drug-target databases to systematically link L2/3-IT-specific differentially expressed genes with existing pharmacological agents. Specifically, detailed drug-target interaction data were first retrieved from the ChEMBL database (Gaulton et al. 2012) and integrated with drug enrichment analysis performed using ToppGene (Chen et al. 2009), in which drug annotations were derived from the Comparative Toxicogenomics Database (CTD) and the STITCH database. Finally, candidate drugs identified from these complementary approaches were cross-compared to identify overlapping compounds consistently supported across different databases and analytical strategies.

### Transcription Factor and Gene Regulatory Network Scoring

To investigate gene regulatory disruptions and restoration in neurons induced by disease and rTMS treatment, the pySCENIC pipeline (v 0.12.1) (Van de Sande et al. 2020) was employed to assess the regulatory activity of transcription factors (TFs). This workflow primarily consists of three steps: (1) construction of co-expression networks using GENIE3, (2) motif enrichment analysis and target gene prediction using RcisTarget, and (3) quantification of regulon activity using AUCell. Regulon specificity scores (RSS) were computed based on cell clusters, and top regulons for each cluster were selected according to the SCENIC protocol implemented in the Python environment. To identify regulons enriched for the U-shaped and inverted U-shaped trajectory genes in neurons, an overlap analysis was performed between these trajectory genes and the gene networks defined by each top regulon. Fisher’s exact test was applied to identify significantly enriched regulons, using a threshold of Benjamini-Hochberg *p*FDR < 0.05. Finally, to visualize the expression patterns of gene networks from the significantly enriched regulons across groups, regulon activity was scored using AUCell and differences between groups were assessed using the rank-sum test with Benjamini-Hochberg *p*_FDR_ < 0.05.

### Cell-cell communication

CellChat (v 1.6.1) (Jin et al. 2021) was employed to infer intercellular communication networks and assess major signaling inputs and outputs across different cell types. The algorithm leverages gene expression profiles and its built-in ligand-receptor interaction database to identify significantly overexpressed ligands and receptors within individual cell populations. Following conversion of cynomolgus monkey genes to their human orthologs, CellChat computed communication probabilities between all pairs of interacting cell populations. Weak interactions were filtered out using default thresholds. Signaling pathway networks were subsequently constructed, enabling the identification and analysis of active signaling pathways among distinct cell types. Finally, the netAnalysis_signalingRole_heatmap function was applied to visualize differences in communication strength across phenotypic conditions using heatmaps.

### Pseudotemporal Analysis of Microglia

Monocle2 (v 2.32.0) (Qiu et al. 2017) was employed to reconstruct the pseudotemporal trajectory of microglia across disease progression, as microglia exhibited the largest number of DEGs in RTT-pre compared to WT. Following conversion of cynomolgus monkey genes to their human orthologs, a CellDataSet object was constructed using exclusively the microglial population. Gene expression data from the three groups were modeled using a generalized linear model (GLM) with a Gaussian family distribution. Significant genes (Benjamini-Hochberg *P*_FDR_ < 0.05) were used as input to the DDRTree algorithm for dimensionality reduction and trajectory reconstruction. The principal cellular trajectory, represented by a minimum spanning tree (MST), was constructed to incorporate all cells into a continuous developmental continuum. Finally, the plot_cell_trajectory function was used to visualize the MST overlaid on cells, allowing observation of cell distribution along pseudotime. To examine the behavior of U-shaped and inverted U-shaped trajectory genes along pseudotime, overlapping genes between these trajectories and the feature genes used in pseudotime analysis were identified. Their expression dynamics were visualized using the plot_genes_in_pseudotime function.

## Data availability

The neuroimaging data supporting the findings of this study are available from the corresponding author upon reasonable request. The snRNA-seq data supporting the findings of this study are available at https://db.cngb.org/search/project/CNP0009605/.

## Acknowledgments

This study was partially supported by STI2030-Major Projects (2021ZD0200900), Natural Science Foundation of China (82125008), National Key Research and Development Program of China (2023YFA1801800), Yunnan Fundamental Research Projects (202501AV070005), Major Basic Research Project of Science and Technology of Yunnan (202505AB350005 and 202102AA100053).

## Compliance with Ethical Standards

All procedures strictly adhered to national guidelines for animal care and use and were approved by the relevant authorities in China and the Institutional Animal Care and Use Committee of Kunming University of Science and Technology (Permit No.: KUST202401002).

## Author contributions

J.W. participated in the study design and significance discussion, provided partial funding support, and offered the experimental platform. Y.C. participated in the study design and significance discussion, provided partial funding support, and supplied model and control monkey resources. T.Y. participated in the study design and significance discussion. W.L. conducted data analysis and manuscript writing. W.S., J.S., and F.Y. participated in experimental operations and contributed to the establishment of the experimental paradigm. X.Q. performed data analysis. Y.Z. managed animal care. Z.Y. provided discussions and suggestions on data analysis. Z.W., Z.C., T.H., H.W., and H.X. carried out animal tissue collection work.

